# Microfluidic Core–Shell Encapsulation Enables Scalable Generation of Apical-Out Intestinal Spheroids

**DOI:** 10.64898/2026.06.10.731294

**Authors:** Pooja Shukla, Ethan J. Taylor, Natsuko Otaki, Kazuki Hattori, Sadao Ota

**Affiliations:** Research Center for Advanced Science and Technology, The University of Tokyo, 4-6-1 Komaba, Meguro-ku, Tokyo 153-8904, Japan; Artificial Intelligence Medicine, Graduate School of Medicine, Chiba University, 1-8-1 Inohana, Chuo-ku, Chiba City, Chiba 260-8670, Japan; Institute for Advanced Academic Research (IAAR), Chiba University, 1-33 Yayoi-Cho, Inage-ku, Chiba City, Chiba 263-8522, Japan; Department of Molecular and Medical Pharmacology, Faculty of Life Sciences, Kumamoto University, 1-1-1 Honjo, Chuo-ku, Kumamoto 860-8556, Japan

**Keywords:** apical-out intestinal spheroids, core-shell microcapsules, high-throughput, microfluidics

## Abstract

Apical-out intestinal spheroids provide direct access to the lumen-facing epithelial surface, making them attractive three-dimensional models for studying epithelial barrier function, nutrient uptake, and luminal exposure. However, existing polarity-reversal methods typically require releasing spheroids from surrounding ECM gels and culturing them in suspension, which can compromise matrix-derived cues, promote fusion, increase size heterogeneity, and limit scalability. Here, we develop a microfluidic core-shell encapsulation strategy to scalably produce apical-out intestinal spheroids within uniform hydrogel microcapsules. These microcapsules consist of a Matrigel core surrounded by an agarose shell. Flow-focusing microfluidics first confines Caco-2 cells in Matrigel cores that provide instructive extracellular matrix cues, and particle-templated emulsification subsequently encloses each core within an inert agarose shell that prevents spheroid fusion and preserves batch uniformity. The method generated >100,000 microcapsules per experiment, with a mean shell diameter of 117 µm, a coefficient of variation below 9%, and >90% single-spheroid formation efficiency. The resulting spheroids established apical–basolateral polarity, organised tight junctions, formed a dextran-excluding epithelial barrier, and exhibited fatty-acid uptake. This core–shell strategy provides an experimentally tractable platform for scalable intestinal epithelial modelling and may be extensible to other epithelial microtissue systems.

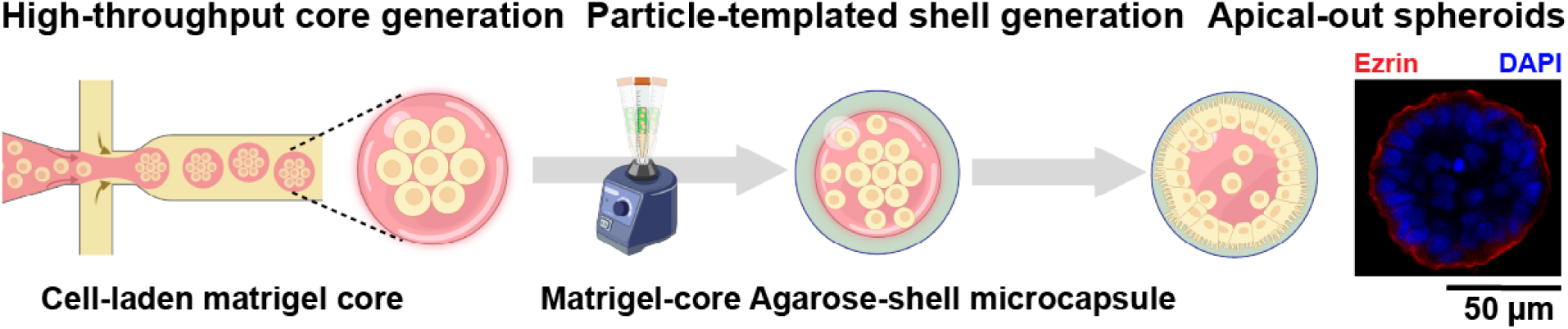

## Introduction

Models of the intestinal epithelium that provide direct access to the lumen-facing surface are essential for studying epithelial transport, barrier functions, and drug absorption. However, generating experimental systems that recapitulate the in vivo physiological dynamics of the intestinal epithelium remains challenging. Although Transwell-based two-dimensional (2D) monolayer cultures are well suited for permeability assays, their physiological relevance for modelling lumen-oriented processes is limited by their inability to recapitulate the three-dimensional (3D) architecture and spatially organised cell–matrix interactions of the native intestinal epithelium (Katt et al., 2016).

Three-dimensional (3D) epithelial models, such as spheroids and organoids, better recapitulate the architecture and functional organisation of the native intestine, including apicobasal polarity and tight junction formation (Co et al., 2019; Kakni et al., 2022; Samy et al., 2019). However, these systems typically adopt an apical-in configuration in which the apical surface is sequestered within a closed lumen. This inward orientation limits direct experimental access to the apical membrane, which is directly exposed to luminal contents, limiting applications such as drug testing, nutrient uptake, and functional studies (Kakni et al., 2022; Wang et al., 2021a).

Apical-out intestinal spheroids have therefore emerged as an attractive alternative in which the apical membrane faces the external environment (Abbas et al., 2023; Ceroni et al., 2025; Co et al., 2019; Nash et al., 2021; Yoshida et al., 2024). This configuration enables direct interrogation of lumen-facing processes, including barrier functions and exposure to xenobiotics. However, current polarity-reversal strategies generally rely on generating spheroids within an extracellular matrix, followed by matrix removal and suspension culture to induce polarity reversal. This process disrupts matrix-derived cues and can result in reduced viability, altered cellular function, and variable outcomes (Csukovich et al., 2024). Moreover, apical-out spheroids are commonly cultured in suspension, diverging from native mechanobiological cues (Wang et al., 2021a). Existing workflows face challenges in both uniformity and scalability. Spheroid size and morphology often vary considerably, generating heterogeneous populations. Furthermore, conventional plate-based methods typically produce only ∼300–1000 spheroids per experiment. Although suspension-based approaches, such as rotating flasks, can increase throughput, they frequently do so at the cost of reduced uniformity and increased operational complexity (Liu et al., 2021).

Here, we present a microfluidic core–shell encapsulation strategy for the scalable generation of apical-out intestinal spheroids. Using a flow-focusing microfluidic device, Caco-2 cells are encapsulated within Matrigel cores that provide spatially confined ECM cues and are subsequently enclosed within inert agarose shells via templated emulsification (Hatori et al., 2018; Maekawa et al., 2026; Otaki et al., 2026). This process yields discrete core–shell microcapsules (mean diameter 117 µm; coefficient of variation 8.72%) that physically isolate individual cell populations without introducing adhesive interactions. Within two days, cells spontaneously self-organise into spheroids that further mature over time within uniform microcapsules, with single-spheroid formation observed in more than 90% of all microcapsules containing cells. The platform enables the production of more than 100,000 spheroids per experiment, overcoming the scalability and heterogeneity limitations of conventional plate-based systems. The resulting spheroids establish apical–basolateral polarity, form intercellular junctions, exhibit epithelial barrier integrity, and support physiologically relevant nutrient uptake. By integrating ECM-guided self-organisation within Matrigel cores with physical confinement imposed by agarose shells, this strategy stabilises outward apical organisation and provides a scalable and experimentally tractable platform for intestinal epithelial modelling and related epithelial microtissue engineering applications.

## Results

### Fabrication of Hybrid core-shell hydrogel microcapsules for intestinal spheroid generation

Epithelial polarity is governed not only by intrinsic cellular programs but also by extrinsic cues, particularly ECM signals that define basal membrane positioning and coordinate tissue organisation (Schweisguth, 2004). Previously, it has been demonstrated in MDCK cells that collagen as ECM induces a basal-out polarity in spheroids, while in suspension culture, devoid of any ECM, cells exhibit apical-out polarity (Wang et al., 1990). Similarly, culturing Caco-2 cells on Matrigel beads have shown to undergo polarity reversal, resulting in apical-out spheroids (Samy et al., 2019). We sought to explore whether this reversal could be achieved in a confined 3D microenvironment. Building on this concept, we hypothesised that encapsulating Caco-2 cells within Matrigel droplets would similarly promote apical-out polarisation within a confined microenvironment. We reasoned that, as the Matrigel polymerises and cells self-organise, spatially restricted ECM cues would be sensed on the interior-facing surface, enabling cells to define their basal domain inward and establish apical identity on the exterior. Matrigel was selected over collagen due to its closer compositional similarity to the native basement membrane and its ability to form stable, soft gels (<0.5 kPa) that recapitulate the mechanical properties of tissues like the small intestine (Cerchiari et al., 2015; Samy et al., 2019).

To test this hypothesis, we encapsulated an average of 15 Caco-2 cells within Matrigel droplets using a flow-focusing device and cultured the beads for one week. Cells self-organised into spheroids, with actin localising to the exterior-facing surface, consistent with apical-out polarity (Supplementary Figure 1a,b). However, these spheroids readily fused over time, resulting in loss of size uniformity and increased heterogeneity within the batch (Supplementary Figure 1a). To address this limitation, we introduced a secondary encapsulation step to physically isolate individual spheroids and prevent fusion. Cell-laden Matrigel beads were encapsulated within a 1% agarose shell via particle-templated emulsification (Hatori et al., 2018), yielding discrete core–shell microcapsules (Figure 1a–c). Agarose, owing to its mechanically inert and non-adhesive nature, enables the physical separation of individual spheroids and thereby preserves size uniformity (Supplementary Figure 1c). Within these microcapsules, Caco-2 cells self-organised into spheroids within two days and underwent further maturation over six days. This approach enabled the consistent generation of more than 100,000 spheroids per experiment (Supplementary Figure 2a,b), demonstrating scalable production.

**Figure 1.**
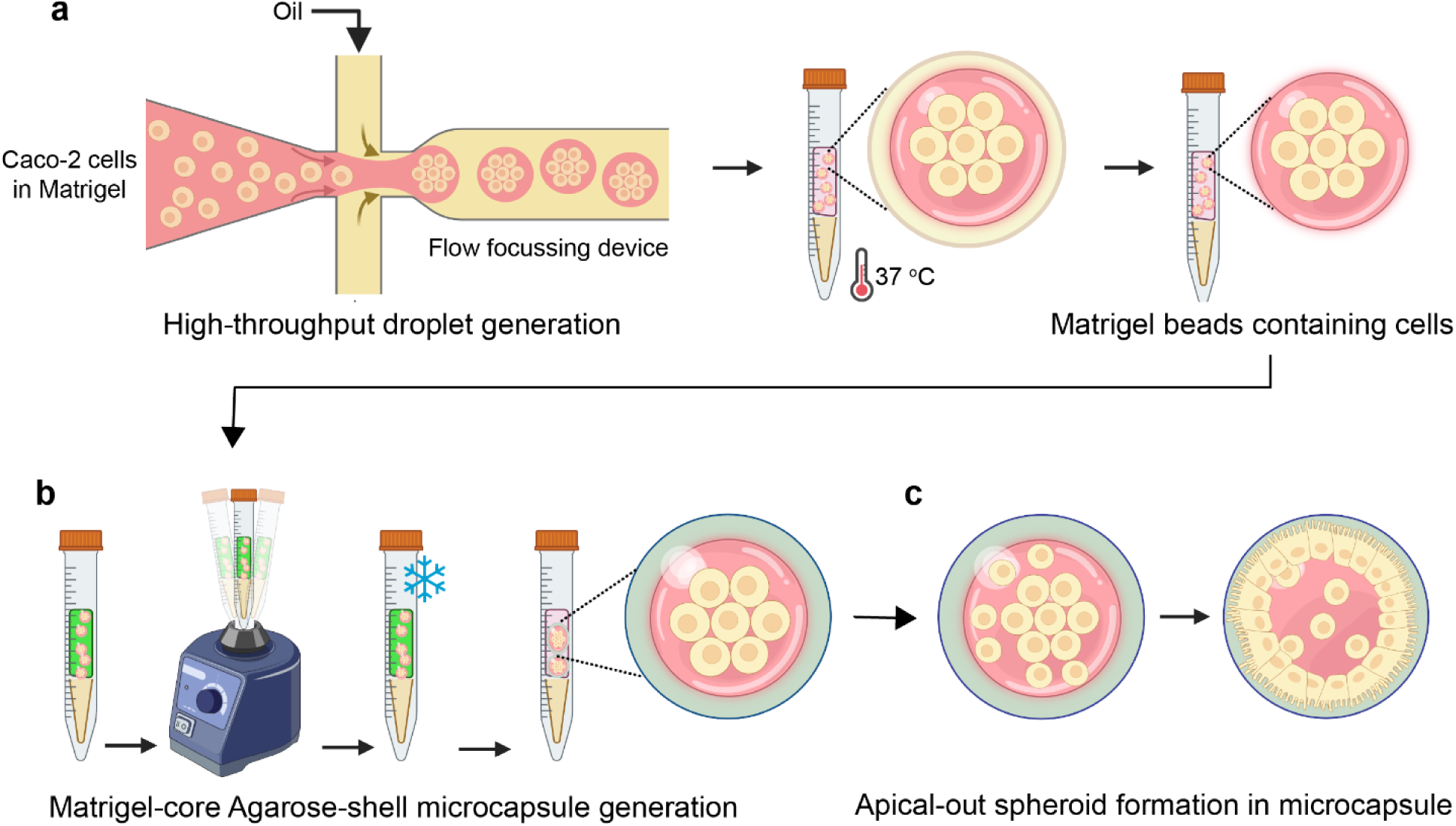
Generation of Hybrid core–shell hydrogel microcapsules for spheroid formation. Schematic showing **a,** Matrigel droplet generation using a flow-focusing microfluidic device. The inner aqueous phase comprises Caco-2 cells suspended in 70% Matrigel, and the outer phase contains fluorinated oil for droplet formation. Droplets were incubated at 37 °C to allow Matrigel polymerisation, and cell-laden Matrigel beads were collected. **b,** the formation of an agarose shell around the Matrigel beads via particle-templated emulsification using a vortex, yielding stable core–shell microcapsules that encapsulate cells within the Matrigel core and **c,** showing self-organisation of encapsulated cells into spheroids inside the microcapsules.

### Hybrid core-shell hydrogel microcapsules enable the formation of intestinal spheroids

To optimise spheroid formation, we systematically evaluated the Matrigel concentration within the microcapsule core. Concentrations of 50%, 70%, and 100% (v/v) were tested, representing low, intermediate, and maximal ECM conditions, respectively. Concentrations below 50% failed to polymerise reliably under microfluidic conditions. Spheroid formation efficiency was quantified at day 2 as the proportion of cell-occupied microcapsules containing a single spheroid-like structure (Figure 2a). Microcapsules with 70% Matrigel exhibited the highest formation efficiency, with 97 ± 3% (mean ± standard deviation across independent experiments) of capsules forming a single spheroid (Figure 2b). A comparable efficiency was observed at 50% Matrigel (93 ± 3%); however, polymerisation at this concentration was inconsistent under loading conditions of 15 cells per bead. In contrast, 100% Matrigel resulted in a reduced formation efficiency of 69 ± 10%, which we attribute to restricted cell mobility within the stiffer matrix, leading to the formation of multiple smaller spheroids within individual microcapsules. Based on these results, 70% Matrigel was selected as the optimal core composition for all subsequent experiments. To assess the overall size uniformity of the microcapsules, we measured their diameters and observed a mean size of 117 μm with a coefficient of variation (CV) of 8.72 %, demonstrating high uniformity and monodispersity across the population. (Figure 2c).

**Figure 2.**
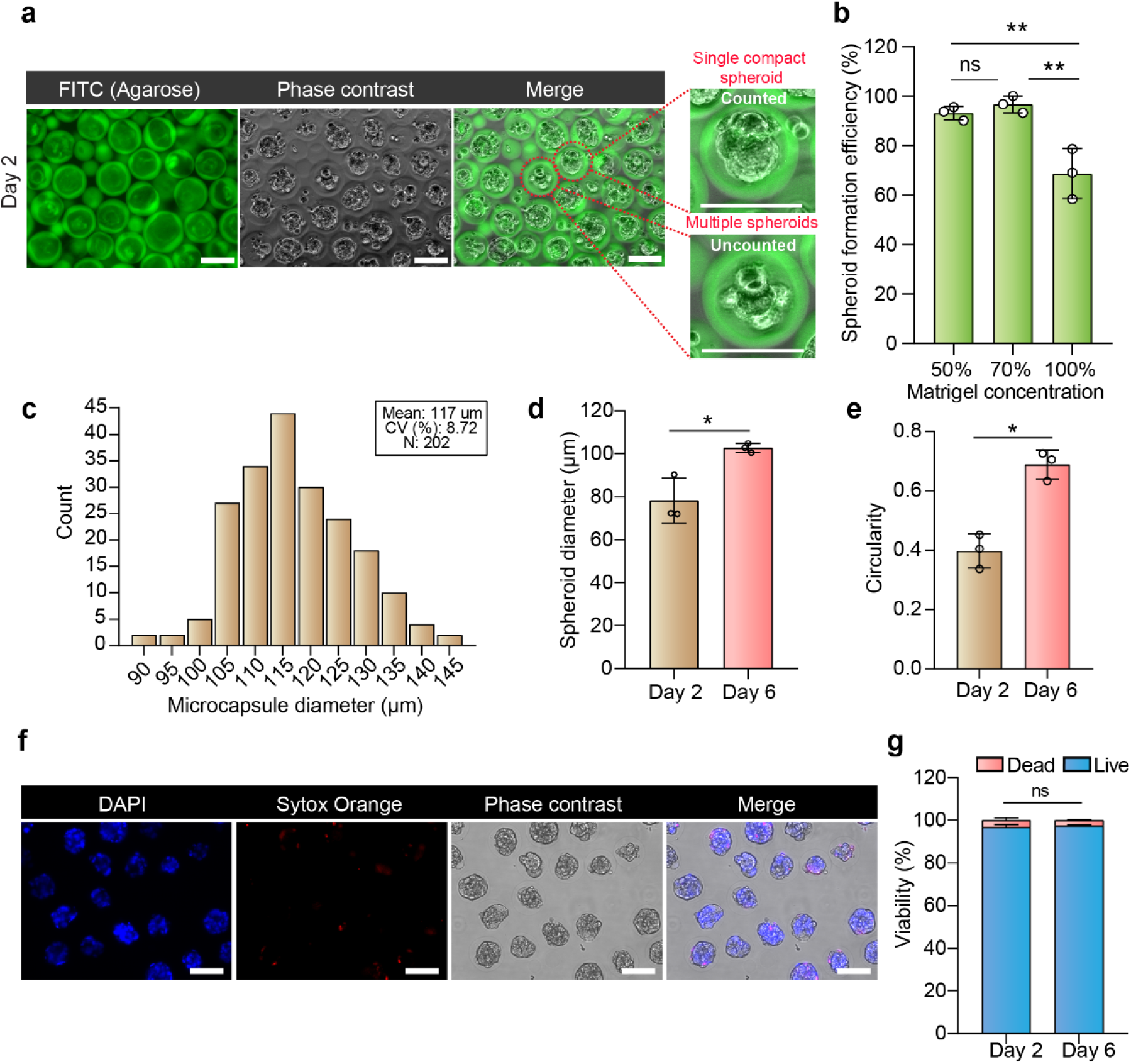
Characterisation of spheroid formation, microcapsule uniformity, growth dynamics, and viability in hybrid core-shell hydrogel microcapsules. **a,** Representative fluorescent and phase contrast images of spheroids in microcapsules. Spheroid formation efficiency was calculated as the fraction of total microcapsules containing a single compact spheroid (top right), while capsules containing multiple spheroids (bottom right) were excluded from the successful category. Scale bars, 100 µm. **b,** Spheroid formation efficiency quantified at day 2 across different Matrigel concentrations, with maximal efficiency observed at 70% Matrigel. Data are shown as mean ± s.d. Statistical analysis was performed using one-way ANOVA (P = 0.0034) followed by Tukey’s multiple comparisons test. ** indicates p-value 0.0080 (50% vs. 100%) and p-value 0.0041 (70% vs. 100%); ns indicates not significant or p-value 0.7830 (50% vs. 70%). **c,** Histogram showing the frequency distribution of microcapsule diameters measured from 202 core–shell microcapsules on day 0. The mean microcapsule diameter is 117 µm with a coefficient of variation (CV) of 8.72%, indicating high size uniformity across the population. Bins represent 5 µm intervals. **d,** Mean spheroid diameter increases from Day 2 to Day 6 in the microcapsules. The bar graph shows the average spheroid diameter per biological replicate measured on day 2 and day 6 (N = 3 replicates, 100 spheroids each). Statistical significance was assessed using an unpaired two-tailed Student’s t-test; * indicates p-value 0.0167 **e,** Circularity of encapsulated spheroids was quantified at day 2 and day 6. Mean circularity increased from 0.39 at day 2 to 0.68 at day 6. The bar graph represents the mean± s.d. of three independent biological replicates (N = 3 replicates, >50 spheroids each), with individual replicate means shown as dots. Statistical significance was determined using a two-tailed paired t-test; * indicates p-value 0.0196. **f,** Representative fluorescence images of spheroids in microcapsules stained with Hoechst (blue) and Sytox Orange (red), alongside phase-contrast and merged views. Images show predominantly Sytox-negative spheroids within agarose-shell microcapsules. Scale bars, 100 µm. **g,** Bar graph showing live and dead cell fractions at day 2 and day 6, assessed by Hoechst/Sytox staining (N = 3 biological replicates, 100 spheroids per replicate). Bars represent mean ± s.d.; the live fraction (Hoechst) is shown in blue and the dead fraction (Sytox) in red. Statistical significance was assessed using an unpaired two-tailed t-test; ns indicates not significant (p -value > 0.05).

We next assessed whether encapsulated spheroids exhibited growth over time. Spheroid diameters were measured at day 2 and day 6, revealing a clear increase from 78.26 ± 12.92 µm to 102.73 ± 18.38 µm (mean ± SD; Fig. 2d and Supplementary Fig. 3a), indicating sustained growth within the microcapsules. In parallel, spheroids underwent marked morphological maturation over time. Quantitative analysis of circularity revealed an increase from a mean value of 0.39 at day 2 to 0.68 at day 6 (Figure 2e), reflecting progressive structural consolidation and acquisition of a more regular epithelial architecture within the confined environment.

To evaluate cellular viability, live/dead staining using Hoechst and SYTOX Orange was performed on day 2 and day 6. Viability remained remarkably high at 98% and showed no significant change over time (Figure 2f,g), indicating that the encapsulated microenvironment supports sustained cellular health during spheroid maturation. Dead cells appeared to be localised to the periphery of the microcapsules and were rarely detected in the spheroid core (Figure 2f). Across all conditions, the dead cell fraction remained below 2%

To assess the temporal stability of the microcapsule system, we quantified agarose shell integrity over an extended culture period of up to 12 days. The proportion of intact shells remained high during the initial culture phase but declined after day 6, reaching 53.3% ± 8.7 (mean ± SD) by day 12 (Supplementary Figure 4a-c). As the majority of spheroid characterisation was conducted at or before day 6, prior to the onset of substantial shell rupture, these analyses correspond to conditions in which the microcapsules remain intact. Within this operational window, no measurable impact of shell integrity on spheroid growth, morphology, or viability was observed, thereby defining the effective stability range of the platform.

### Spheroids display apical-out polarity inside hybrid core-shell hydrogel microcapsules

It is well established that cortical actin delineates the apical domain in polarised epithelial cells, providing a structural framework for apical surface organisation. In parallel, ezrin, a canonical brush border marker and member of the ERM (ezrin–radixin–moesin) family, links actin filaments to the apical plasma membrane and is critical for stabilising microvillar architecture (Pelaseyed and Bretscher, 2018). To check the establishment of epithelial polarity within the core–shell microcapsules, we performed immunofluorescence staining for actin and ezrin in day 6 spheroids. Staining revealed the formation of a continuous Actin ring on the outer surface of the spheroids. In addition, ezrin staining showed strong enrichment at the same apical surface, where it colocalised with actin, defining the apical domain of the spheroids (Figure 3a).

**Figure 3.**
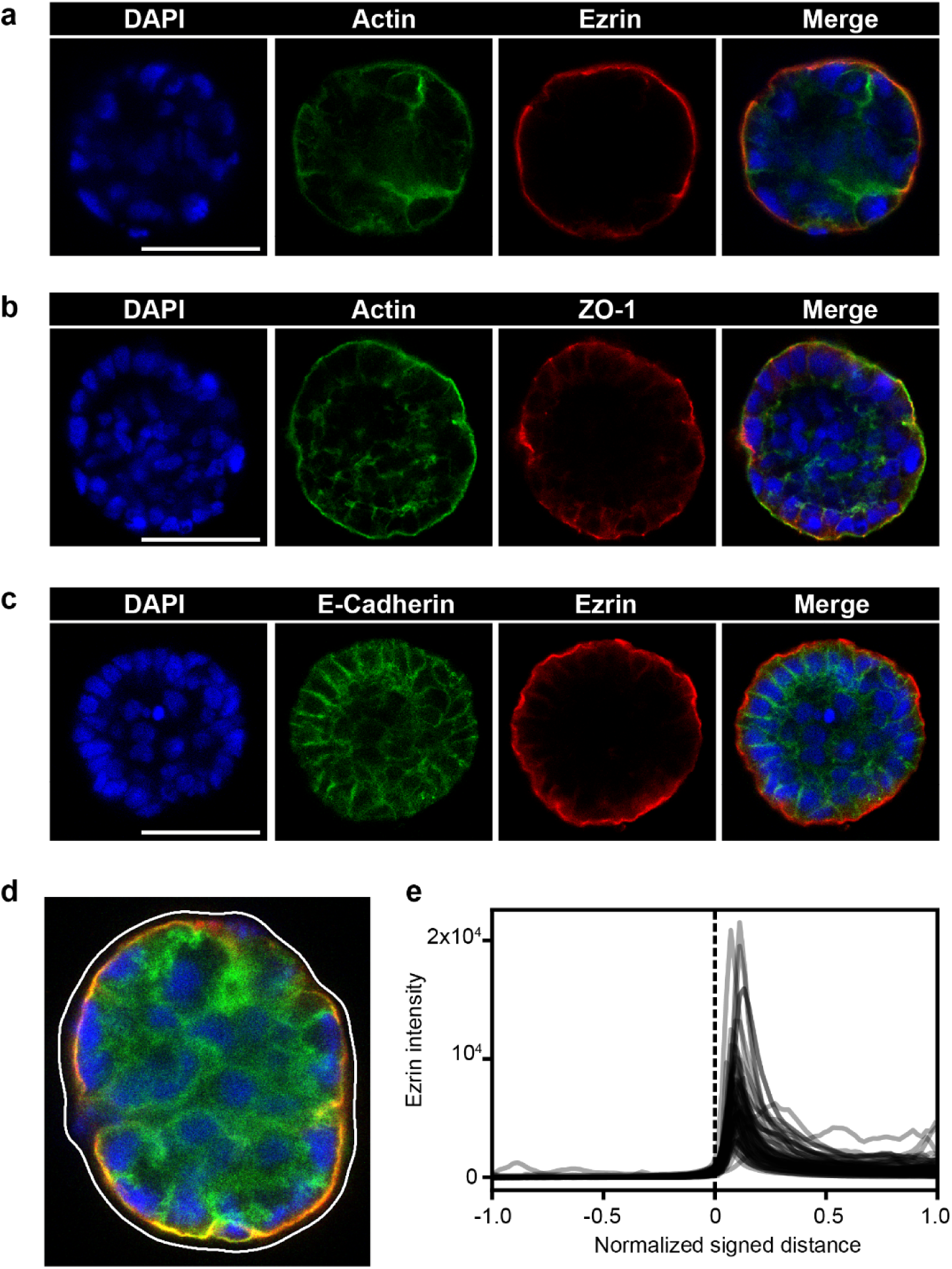
Immunofluorescence analysis of epithelial polarisation in day 6 spheroids. **a,** Immunofluorescence staining for Ezrin (red) and actin (green) highlights apical polarisation of intestinal spheroids within core–shell microcapsules. Actin enrichment at the apical domain colocalises with Ezrin, indicating the establishment of apical membrane identity. Nuclei are counterstained with DAPI (blue). Scale bar, 50 µm. **b,** Immunostaining for the tight junction protein ZO-1 (red) showing ring-like distribution at cell–cell interfaces, colocalising with actin (green) and confirming the formation of functional tight junctions. Nuclei are counterstained with DAPI (blue). Scale bar, 50 µm. **c,** Immunofluorescence staining for E-cadherin (green) and Ezrin (red) demonstrates spatial segregation of basolateral and apical membrane domains within encapsulated spheroids. E-cadherin localises predominantly to lateral cell–cell junctions, forming a continuous adherens junction network. This complementary distribution confirms the establishment of apical–basolateral polarity. Nuclei are counterstained with DAPI (blue). Scale bar, 50 µm. **d,** Representative confocal image showing Ezrin, Actin, and DAPI immunofluorescence staining. The spheroid boundary obtained from the combined segmentation of actin and DAPI signals is overlaid in white. **e,** Radial intensity profiles of ezrin as a function of normalised signed distance from the spheroid boundary. Each grey trace represents an individual spheroid, and >50 spheroids across three biological replicates were analysed. Distance is normalised from −1 (outside) to +1 (centre), with 0 indicating the spheroid boundary (dashed line). Ezrin intensity peaks sharply at the boundary and decreases toward the interior, indicating preferential enrichment at the outer surface.

Junctional complexes are known to associate with the actin cytoskeleton, and their maturation requires actin reorganisation. Among them, ZO-1 functions as a key scaffold protein that links claudins to the actin network(Fanning et al., 2012; Hartsock and Nelson, 2008). Consistent with this, our staining revealed clear colocalisation of ZO-1 with cortical actin at the spheroid surface, indicating the coordinated assembly of tight junctions with the apical actin cytoskeleton (Figure 3b). E-cadherin displayed a lateral distribution along cell–cell interfaces, forming a continuous circumferential network consistent with adherens junction assembly and epithelial cohesion. The spatial segregation of ezrin to the outer membrane and E-cadherin to lateral junctions reveals the emergence of apical–basolateral polarity within the microcapsules (Figure 3c).

Next, to quantitatively assess apical polarity, we analysed the spatial distribution of ezrin using confocal microscopy. Spheroids were segmented using a composite actin and DAPI signal to define spheroid boundaries (Figure 3d). Ezrin localisation was then quantified relative to the boundary using normalised radial coordinates, enabling direct comparison across spheroids of different sizes. Radial intensity profiles revealed a pronounced peak in ezrin signal at the spheroid boundary, followed by a progressive decline towards the interior (Figure 3e). This peripheral enrichment was consistently observed across more than 50 spheroids analysed from three independent experiments, indicating robust localisation of ezrin to the outer surface and confirming apical-out polarity.

Together, these results demonstrate that the core–shell microcapsule system supports the formation of structurally organised and polarised intestinal epithelial spheroids with well-defined apical-out domains and functional intercellular junctions.

### Epithelial barrier formation in spheroids within microcapsules

A principal function of epithelial tissues is to establish a selective barrier that restricts paracellular permeability. This barrier is mediated by apical tight junctions, which form intercellular seals between neighbouring cells. To determine functional barrier integrity, we used 70 kDa FITC–dextran to selectively probe unrestricted permeability, as it is excluded from tight junction–mediated transport (Horowitz et al., 2023). Spheroids at day 6 were subjected to a FITC–dextran (70 kDa) diffusion assay. Under control conditions, spheroids effectively excluded FITC–dextran from the interior, with fluorescence largely confined to the external capsule environment (Figure 4a). Disruption of calcium-dependent adhesion with EGTA (10 mM) abrogated this barrier function, as FITC–dextran penetrated into the spheroid core, accompanied by loss of epithelial organisation (Figure 4a).

**Figure 4.**
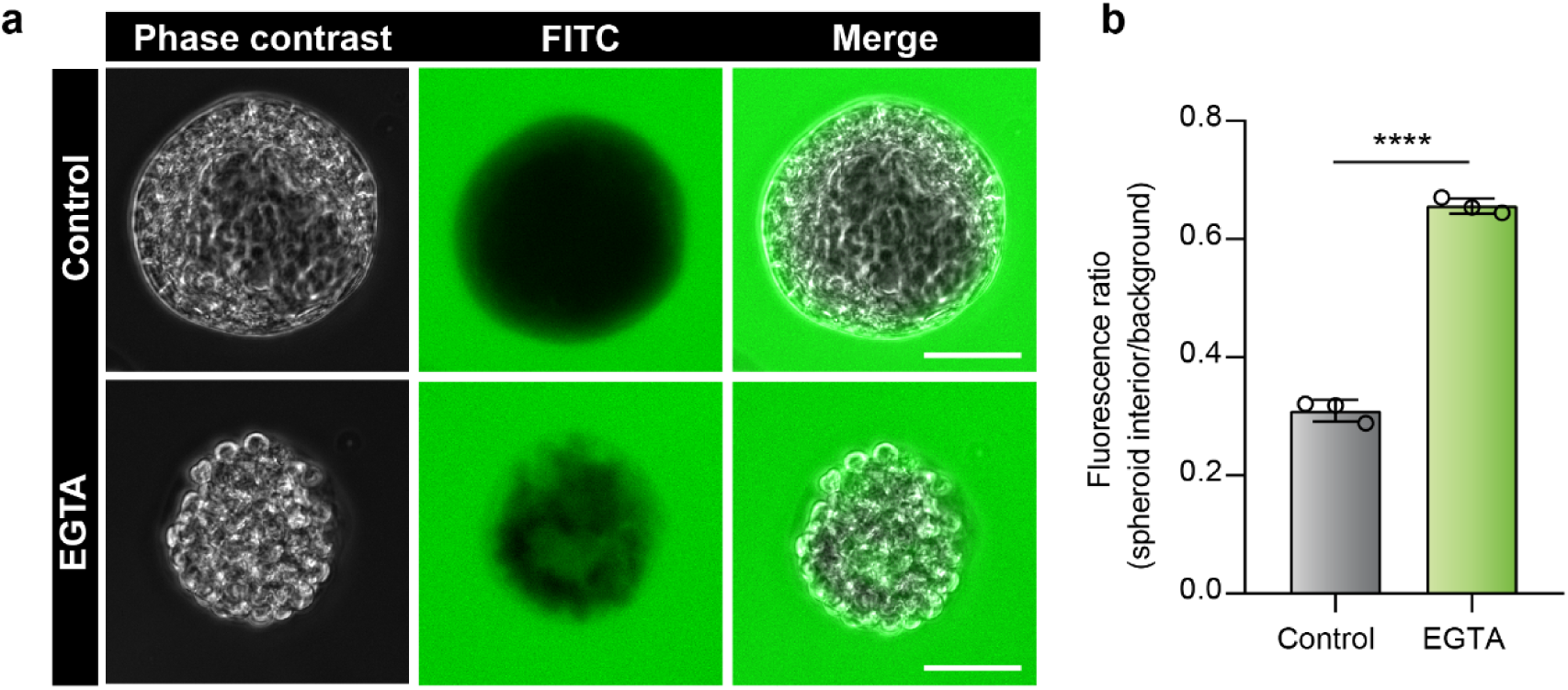
Functional assessment of barrier integrity in intestinal spheroids. **a,** FITC–dextran permeability assay on day 6 showing intact spheroids excluding FITC–dextran (top) and EGTA-treated spheroids displaying FITC penetration into the spheroid (bottom), indicating compromised barrier integrity. Scale bars, 50 µm. **b,** Quantification of FITC–dextran permeability expressed as the fluorescence intensity ratio between the spheroid interior and background. EGTA treatment significantly increased dextran penetration compared to control spheroids. Bars represent mean ± s.d. from three independent biological replicates (N = 3 replicates, >50 spheroids each), with individual replicate means shown as dots. Statistical significance was determined using a two-tailed unpaired t-test; **** indicates p-value <0.0001.

Barrier function was quantified by measuring the fluorescence intensity ratio between the spheroid interior and the surrounding background. Control spheroids displayed a low interior-to-background ratio (0.30), consistent with effective exclusion of dextran. In contrast, EGTA-treated spheroids showed a significantly elevated ratio (0.65), reflecting increased permeability (Figure 4b; Supplementary Figure 5a). Together, these results demonstrate that the impermeability of untreated spheroids is dependent on calcium-mediated junctional integrity and confirm the functional maturation of epithelial barriers within the core–shell microcapsule system.

### Functional nutrient uptake across polarised intestinal spheroids

To determine whether apical-out intestinal spheroids exhibit functional nutrient uptake and directional transport, we assessed trafficking of the fluorescent fatty acid analogue C1-BODIPY-C12 (Co et al., 2019; Kakni et al., 2023a). The probe was added to the extracellular medium, and uptake was analysed by fluorescence microscopy. In apical-out spheroids, C1-BODIPY-C12 accumulated predominantly at the basolateral surface, consistent with apical-to-basal transport across the epithelial layer (Figure 5a). As a control, suspension Caco-2 spheroids (Kakni et al., 2023b; Wang et al., 2021b) were analysed in parallel. Quantification of intracellular fluorescence revealed no significant differences between spheroids in microcapsules and spheroids in suspension, indicating that spheroids generated in the core–shell microcapsule system recapitulate the functional performance of established models (Figure 5b; Supplementary Figure 6a). These findings demonstrate that spheroids formed within the Matrigel core–agarose shell not only acquire structural polarity but also display physiologically relevant absorptive function.

**Figure 5.**
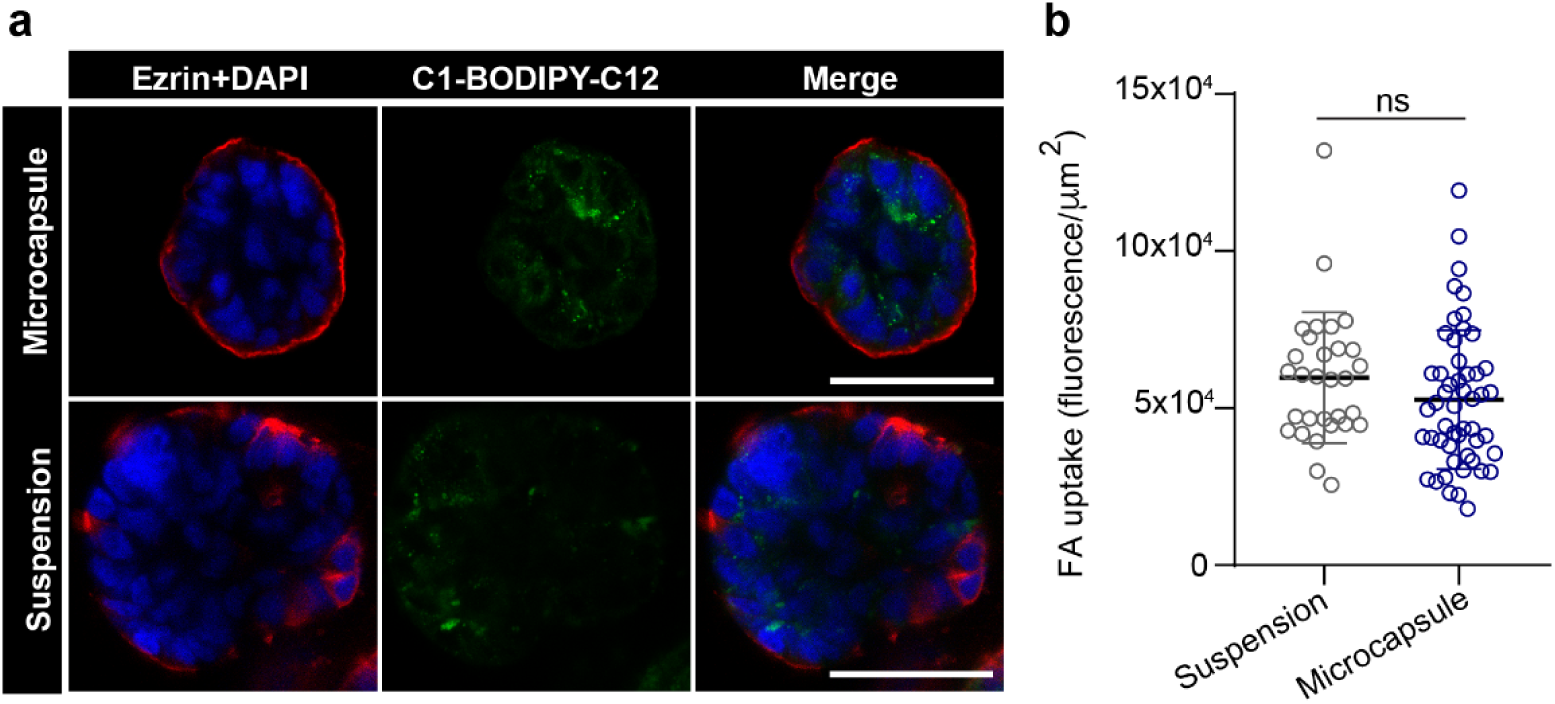
Comparative analysis of fluorescent fatty acid uptake in day 6 spheroids generated by suspension culture versus microcapsules. a, Confocal imaging of C1-BODIPY-C12 uptake shows comparable fatty acid incorporation in spheroids formed within microcapsules and those generated by suspension culture. Scale bars, 50 µm. b, Quantification of fatty acid analogue (FA) uptake across spheroids demonstrates no significant difference between conditions. Data are presented as mean ± s.d. (*n* = 50 spheroids). Statistical analysis was performed using the Mann–Whitney *U* test; ns indicates not significant (p-value 0.1159).

## Discussion

The hybrid core–shell hydrogel microcapsule system presented here offers several key advantages for modelling epithelial polarity and function. The use of Matrigel as the ECM core provides physiologically relevant biochemical and mechanical cues, while the agarose shell serves as an inert, non-adhesive barrier that prevents spheroid fusion and preserves uniformity. By integrating microfluidic droplet generation with particle-templated emulsification, the platform achieves high-throughput encapsulation with excellent monodispersity and scalability. Optimisation experiments identified 70% Matrigel as the optimal core composition, as it not only overcomes the polymerisation inconsistencies observed at 50% but also provides sustained ECM cues to support epithelial organisation over a week-long culture. The platform reliably enables production of over 100,000 intestinal spheroids in microcapsules within 3 hours of experimental time and yields mature apical-out spheroids within 6–7 days, demonstrating exceptional throughput, reproducibility, and compatibility with standard culture workflows. The resulting microcapsules are compact in size (mean diameter 117 μm), which simplifies handling and facilitates downstream processing. Furthermore, the relatively small size and optical transparency of the capsules may help limit light scattering, facilitating faster and higher-resolution 3D imaging across large numbers of spheroids and thus supporting high-throughput phenotypic analysis.

Despite these strengths, the approach also presents certain limitations. Handling Matrigel within a microfluidic setup remains challenging due to its temperature sensitivity and tendency to polymerise during droplet generation, which can disturb encapsulation efficiency and lead to material loss. This is particularly evident at higher input volumes (>100 μL), and we recommend restricting sample inputs to ≤100 μL to minimise waste. Incomplete polymerisation of Matrigel droplets can further result in occasional sample loss. Additionally, because Matrigel is mechanically soft, the encapsulated cores may not always remain perfectly spherical during agarose shell formation, sometimes yielding irregular morphologies or even partial disintegration of beads during encapsulation. For long-term cultures (>10 days), spheroids can occasionally outgrow the confining shell, leading to breaches or fusion events. We therefore recommend culture and analysis within 6–7 days, when apical-out polarity and barrier formation are robustly established. Future studies will be needed to evaluate the stability and functional maturation of spheroids in this system under extended culture conditions (>15 days).

The 1% agarose shell allows free diffusion of nutrients and growth factors, as evidenced by consistently high cell viability and the presence of only rare peripheral dead cells (<5%) (Figure 2, f,g; supplementary figure 2c). Importantly, the agarose shell can be enzymatically dissolved using agarase, permitting the recovery of intact spheroids for downstream use (Otaki et al., 2026). This feature may enhance the versatility of the system for both basic research and potential translational applications.

The modularity of the core–shell design suggests potential application beyond intestinal models. On one hand, by tuning ECM composition, the system could be adapted to generate apical-out spheroids from other epithelial tissues, such as lung or nasal epithelia, or to support organoid cultures where Matrigel is essential for differentiation. On the other hand, its barrier-forming capability suggests potential utility in modelling complex interfaces such as the blood–brain barrier or placental barrier. The compartmentalised architecture is also compatible with high-throughput analytical workflows: microcapsules can be sorted individually based on fluorescence or phenotypic markers using flow cytometry, enabling selection of spheroids according to functional behaviour. This aligns well with current demands in drug screening and precision medicine.

In summary, the Matrigel–core agarose–shell microcapsule system provides a scalable and versatile platform for generating uniform, functional, apical-out intestinal spheroids. By combining structural polarity, barrier integrity, and nutrient uptake with high-throughput scalability, the method overcomes key limitations of existing approaches and establishes a framework adaptable to diverse applications in basic research, disease modelling, and biotechnology.

## Materials and Methods

### Cell culture

Caco-2 cells (ATCC, HTB-37) were cultured in DMEM-high glucose (Wako, 045-30285) containing 10% (v/v) heat-inactivated fetal bovine serum (Sigma, F7524), 2 mM L-alanyl-L-glutamine (Wako, 016-21841), 1X nonessential amino acids (Sigma, M7145), and 1X Antibiotic-Antimycotic (Thermo Fisher Scientific, 15240096) in a humidified atmosphere with 5% CO2 at 37°C. For passaging, cells were briefly treated with TrypLE (Gibco,12604013). Cells were regularly tested negative for mycoplasma using Cycleave PCR Mycoplasma Detection Kit (Takara Bio, CY232).

### Preparation of FITC-Agarose

Ultra-Low Gelling Temperature Agarose (Sigma, A5030-5G) was dissolved in dimethyl sulfoxide (DMSO; Wako, 048-32811) at 25 mg/mL, vortexed, sealed with a septum, and bubbled with nitrogen. The solution was heated at 85 °C for 4–5 h to evaporate DMSO. Fluorescein isothiocyanate isomer-I (FITC; Wako, F007) was dissolved in DMSO (100 mg/mL) and mixed 1:1 with the agarose pellet, followed by stirring at 40 °C for 1 h. The FITC–agarose was lyophilised, reconstituted in Milli-Q water (1% w/v), and dissolved at 55 °C. Solutions (1 mL) were centrifuged at 20,000 × g, 37 °C for 3 min, and the supernatant was re-centrifuged 2–3 times until clear. FITC–agarose aliquots were stored at 4 °C. For the templated emulsification step (in Fig. 1), A 2% agarose solution was prepared in PBS and mixed with 1% FITC–agarose at 1:9 (v/v).

### Generation of intestinal spheroids in Matrigel-core Agarose-shell microcapsules

Cell preparation for encapsulation: Cells at ∼80% confluence in 10-cm dishes (Corning, 430167) were trypsinised and collected. The suspension was treated with 20 µg/mL deoxyribonuclease I (Wako, 047-26771) in PBS and gently agitated for 15 min at room temperature to minimise aggregation. From this step onward, all procedures were performed on ice or at 4°C. Cells were washed once with cold culture medium and centrifuged at 200 × g for 3 min at 4 °C. After counting, ∼5.6 × 10⁶ cells were prepared, corresponding to the Poisson distribution for λ = 15 in 80-µm droplets. The cell pellet was resuspended in 100 µL of 70% (v/v) Matrigel in culture medium.

Microfluidics device fabrication: Flow-focusing devices were fabricated by standard soft lithography using polydimethylsiloxane (PDMS; SILPOT184, DOW). Each device comprised two inlets—one for the cell suspension and one for Automated Droplet Generation Oil for EvaGreen (Bio-Rad, 1864112)—and a single outlet.

Generation of cell-laden Matrigel cores: The microfluidic setup was maintained at 4 °C in a cold chamber to prevent premature Matrigel polymerisation. Cell suspensions were introduced into the channels via PEEK tubing using FLPG Plus and LineUp Flow EZ controllers (Fluigent, LU-FEZ-2000) at 350 mbar, while Droplet Generation Oil was supplied by a syringe pump (Harvard Apparatus, PUMP 11 Elite) at 25 µL/min. Droplets were collected in microcentrifuge tubes (Eppendorf, 0030123620) and incubated at 37 °C for 30 min, with gentle tapping every 10 min to ensure uniform polymerisation. The emulsion was then broken using 20% (v/v) 1H,1H,2H,2H-perfluoro-1-octanol (PFO; Wako, 324-90642) in HFE-7200 (3M), and the Matrigel beads were collected in culture medium. The emulsion-breaking step was repeated 2–3 times to remove any residual oil and unpolymerized aggregates.

Core-shell microcapsule generation: Cell-laden Matrigel beads were pelleted at 100 × g for 3 min in 15-mL tubes (Eppendorf, 0030122259), and the supernatant was removed. An equal volume of 2% (v/v) Ultra-Low Gelling Temperature Agarose (Sigma, A5030-5G) in PBS was added and gently mixed. Automated Droplet Generation Oil for EvaGreen (Bio-Rad, 1864112), at five times the bead/agarose volume, was then added, and the tube was vortexed at maximum speed for 30 s to achieve emulsification. The emulsion was incubated on ice for 10 min to solidify the agarose shell, forming two-layered Matrigel–agarose gels with cells encapsulated in the core. Emulsions were broken using 20% (v/v) 1H,1H,2H,2H-perfluoro-1-octanol (PFO; Wako, 324-90642) in HFE-7200 (3M), and the microcapsules were collected in culture medium. This step was repeated 2–3 times to remove residual oil and unpolymerized aggregates. The resulting Matrigel-core agarose-shell microcapsules were cultured in ultra-low attachment plates (Corning, 3471) with cell culture medium.

### Cell viability assay and analysis

Spheroids were harvested on day 2 and day 6, washed once in phosphate-buffered saline (PBS), and resuspended in a staining solution consisting of 1 μg/mL Hoechst 33342 (Dojindo, 346-07951) and 0.25 μM SYTOX™ Orange Nucleic Acid Stain (Thermo Fisher Scientific, S32861) in PBS. The suspension was incubated at 37 °C for 15 minutes in the dark, after which the microcapsule concentration was adjusted to achieve optimal imaging density. Samples were imaged immediately using the EVOS FL Auto Imaging System (Thermo Fisher Scientific, C10228) with a 20× air objective.

Image analysis was performed using Fiji/ImageJ (version 1.54). Regions of interest (ROIs) corresponding to individual spheroids were manually defined using phase-contrast images and subsequently applied to the corresponding Hoechst and SYTOX Orange fluorescence channels. Each fluorescence image was converted to 8-bit grayscale and thresholded using Otsu’s method, followed by binary conversion. The integrated pixel area within each ROI, representing the fluorescence signal from either live (Hoechst-positive) or dead (SYTOX-positive) cells, was quantified using the *Measure* function.

For each spheroid, the live and dead fractions were calculated as the ratio of the respective signal area to the total signal area (i.e., Hoechst + SYTOX). The percentage of dead cells was defined as: Dead fraction (%) = SYTOX area / (Hoechst area + SYTOX area) × 100. These values were analysed per spheroid.

### Spheroid size analysis

Spheroid size was quantified from brightfield images using Fiji/ImageJ (version 1.54). Images were first converted to 8-bit grayscale and thresholded using Otsu’s method to generate binary masks of individual spheroids. Morphological operations, i.e., closing and hole filling, were applied to obtain continuous spheroid outlines. Regions of interest (ROIs) corresponding to individual spheroids were manually selected, and size measurements were obtained using the *Measure* function. Spheroid size was defined as the Feret diameter (maximum calliper diameter) of each ROI.

### Circularity analysis

Spheroid circularity was quantified from brightfield images using Fiji/ImageJ (version 1.54). Images were first converted to 8-bit grayscale and thresholded using Otsu’s method to generate binary masks of individual spheroids. Morphological operations, i.e., closing and hole filling, were applied to obtain continuous spheroid outlines. Regions of interest (ROIs) corresponding to individual spheroids were manually selected and analysed using the *Measure* function. Circularity was calculated as 4π × area/perimeter^2^, with values ranging from 0 (irregular shape) to 1 (perfect circle).

### Immunofluorescence staining of spheroids

Spheroids were fixed in 4% formaldehyde (in PBS) for ≥15 min at room temperature, washed three times with PBS, and permeabilised with 0.5% Triton X-100 (v/v) in 1% BSA for 10 min. After PBS washing, samples were blocked with 5% BSA for 1 h at room temperature. Primary antibodies, diluted in 2% BSA at manufacturer-recommended concentrations, were incubated overnight at 4 °C with gentle agitation. Following three PBS washes, Alexa Fluor–conjugated secondary antibodies (1:500 in 2% BSA) were applied for 2 h at room temperature. Nuclei were counterstained with DAPI for 5 min at room temperature. Spheroids were mounted on glass slides using ProLong™ Glass Antifade Mountant (Invitrogen, P36982).

The following antibodies were used in the study: Actin (Phalloidin-iFluor™ 488 Conjugate; Cayman, 20549), Ezrin polyclonal antibody (Proteintech, 26056-1-AP), ZO-1 polyclonal antibody (Proteintech, 1773-1-AP), E-cadherin polyclonal antibody (BD Biosciences, 610181), Goat Anti-Rabbit IgG H&L (Alexa Fluor® 647) preadsorbed (abcam, ab150087), Goat anti-mouse AF 488 (abcam, ab150113).

Samples were imaged using a Leica Stellaris confocal microscope with a 63× oil-immersion objective or EVOS FL Auto Imaging System (Thermo Fisher Scientific, C10228) with a 20× air objective. Representative fields were acquired from each sample and pseudo-colours were assigned according to the respective secondary antibodies and dyes.

### Ezrin enrichment analysis

Confocal spheroid images stored as Leica ‘.lif’ files were analyzed in Python using liffile, NumPy, SciPy, and scikit-image. Only 2D cross-sectional images located at the central thickness of the spheroid containing DAPI, actin, and ezrin channels were used for quantitative analysis. Spheroid segmentation was performed from the DAPI and actin channels, which were independently percentile-normalised (1st–99.5th percentiles), summed, and Gaussian-smoothed with a kernel size scaled to image dimensions. The combined signal was then globally thresholded using Li’s method. The resulting binary mask was refined by morphological closing, hole filling, and removal of small objects, and connected components were labeled as individual spheroids. Spheroid boundaries were obtained from the final binary mask as the one-pixel inner boundary generated by mask erosion; the longest contour was extracted for visualization. Objects touching the image border were excluded from downstream quantification to ensure only whole spheroids were analyzed.

Ezrin localisation was quantified relative to the spheroid boundary. A signed distance map was computed for each segmented spheroid, with pixels inside the spheroid assigned positive distances and pixels outside assigned negative distances. Distances were normalised separately by the maximum interior and exterior distances to generate a dimensionless radial coordinate ranging from −1 to 1, where 0 corresponds to the spheroid boundary. Radial ezrin intensity profiles were generated by binning fluorescence intensity values as a function of normalised boundary distance. The resulting profiles enabled quantitative comparison of ezrin distribution across spheroids of different sizes and were used to assess preferential localisation of ezrin at the spheroid periphery.

### Barrier integrity Assay

Spheroids were washed with PBS, pelleted at 100 × g for 3 min, and resuspended in either 1 mg/ml 70 kDa FITC–dextran (Sigma-Aldrich, 46945-100MG-F) or 1 mg/ml 70 kDa FITC–dextran with 10 mM EGTA (Wako, QB-6401). Samples were incubated for 15 min at 37 °C in the dark. The spheroids were mounted on slides, and density was adjusted for optimal imaging. Fluorescence intensity within the spheroid interior was quantified immediately after imaging.

Image analysis was performed using Fiji/ImageJ (version 1.54). Phase-contrast images were used to define spheroid geometry independently of fluorescence intensity. Spheroids were segmented by thresholding followed by morphological opening and closing and hole filling, and individual spheroid boundaries were identified using particle analysis. To exclude surface-associated and optical edge effects, spheroid masks were uniformly eroded by 10 µm, and the resulting inner regions were re-segmented to generate per-spheroid inner ROIs. These ROIs were applied to the corresponding raw 16-bit FITC fluorescence images, and mean fluorescence intensity was quantified per spheroid. Background fluorescence was measured from ROIs placed in spheroid-free regions of the same image. All measurements were performed on uncompressed TIFF images using identical acquisition and analysis parameters across conditions.

### Fatty acid uptake assay

Spheroids were washed with phenol red–free DMEM (FluoroBrite™ DMEM, A1896701) and resuspended in 5 μM fluorescent fatty acid analogue C1-BODIPY-C12 (Invitrogen, D3823) supplemented with 5 μM fatty acid–free BSA (Wako, 9048-46-8). Spheroids were plated in low-attachment 24-well plates, incubated for 30 min, and fixed in 4% paraformaldehyde in PBS. Following nuclear and actin staining, single optical sections through the central plane of each spheroid were acquired by confocal microscopy.

Fatty acid uptake was quantified in Fiji/ImageJ (version 1.54). Spheroid boundaries were manually delineated from the actin channel and applied as regions of interest (ROIs) to the corresponding C1-BODIPY-C12 channel. For each spheroid, the area and raw integrated fluorescence density (Raw IntDen) were measured. Mean background fluorescence was determined from three cell-free regions within the same image and used to calculate background-corrected integrated density according to: Corrected IntDen = Raw IntDen − (Background Mean × Number of Pixels)

Fatty acid uptake was expressed as the background-corrected integrated density normalised to spheroid area (fluorescence/µm²). More than 50 spheroids from three independent experiments were analysed for each condition. For control (suspension) spheroids, microcapsules were dissolved in 5 mM EDTA in PBS on a rotating platform at 4 °C for 1 h prior to incubation with C1-BODIPY-C12 under identical conditions. For negative controls, spheroids were incubated with vehicle (DMSO) instead of C1-BODIPY-C12 while maintaining all other conditions unchanged.

### Shell Integrity assessment

Shell stability was assessed by quantifying the proportion of intact microcapsules over time. Agarose shells were labelled with FITC to visualise shell continuity. Approximately 100 microcapsules were seeded per well in a 96-well plate (day 0) and imaged in their entirety at days 0, 3, 6, 7, 8, 9, 10, 11 and 12. Day 6 represented the standard endpoint, and culture was extended to day 12 to evaluate long-term stability. Experiments were performed in three independent biological replicates. An intact microcapsule was defined as a single spheroid enclosed within a continuous, unbroken FITC-labelled agarose ring. Capsules showing rupture, discontinuity of the fluorescent shell, or spheroid extrusion were classified as non-intact.

Shell integrity (%) was calculated as: Intact shells (%)= (Number of intact microcapsules at time t/Total microcapsules seeded at day 0) ×100

### Statistical analysis

All statistical analyses were performed using GraphPad Prism version 8. Comparisons between two groups were conducted using paired or unpaired two-tailed *t* tests, as appropriate. For comparisons among three or more groups, one-way ANOVA was applied. The number of biological replicates (N), sample sizes (n), and the statistical tests used are specified in the corresponding figure legends. Significance was defined as *p* < 0.05, with *p* > 0.05 considered not significant (ns). Exact *p-*values are reported in the figure legends.

### Data availability

The analysis code supporting this study is available at solabtokyo-org/spheroid-enrichment

## Author contributions

Conceptualisation, P.S. and S.O.; Methodology, P.S.; Validation, P.S.; Formal Analysis, P.S.; E.T. (Ezrin enrichment analysis); Investigation, P.S.; Resources, P.S. and S.O.; Data Curation, P.S.; Writing – Original Draft, P.S. and S.O.; Writing –Review & Editing, P.S., N.O., E.T., K.H., and S.O.; Supervision, S.O.; Funding Acquisition, N.O., K.H., and S.O.

## Acknowledgements

The authors express their sincere gratitude to all members of the Ota Laboratory at RCAST, The University of Tokyo, for their constructive discussions and support. Figure 1 and Supplementary Figure S1a were created using BioRender.com.

This work was supported by multiple funding sources. We gratefully acknowledge the Japan Science and Technology Agency (JST) through the Core Research for Evolutionary Science and Technology (CREST) program (JPMJCR23B6 to S.O.), the Adopting Sustainable Partnerships for Innovative Research Ecosystem (ASPIRE) program (JPMJAP2416 to S.O.), the Precursory Research for Embryonic Science and Technology (PRESTO) program (JPMJPR24N3 to N.O.), the Fusion Oriented Research for Disruptive Science and Technology (FOREST) program (JPMJFR240K to K.H.), and the GteX program (JPMJGX23B1 to S.O.).

Support was also provided by the Japan Agency for Medical Research and Development (AMED) under grant numbers JP22gm6710008 (to K.H.) and JP256f0137008 (to N.O.).

The authors further acknowledge support from the Japan Society for the Promotion of Science (JSPS KAKENHI) through grants JP22K12797 (to K.H.), JP25K18827 (to N.O.) and 25H01359 (to S.O.). Additional support was provided by the Takeda Science Foundation Life Science Research Grant and the Nakatani Foundation (to S.O.).

## Supplementary

**Supplementary Figure 1.**
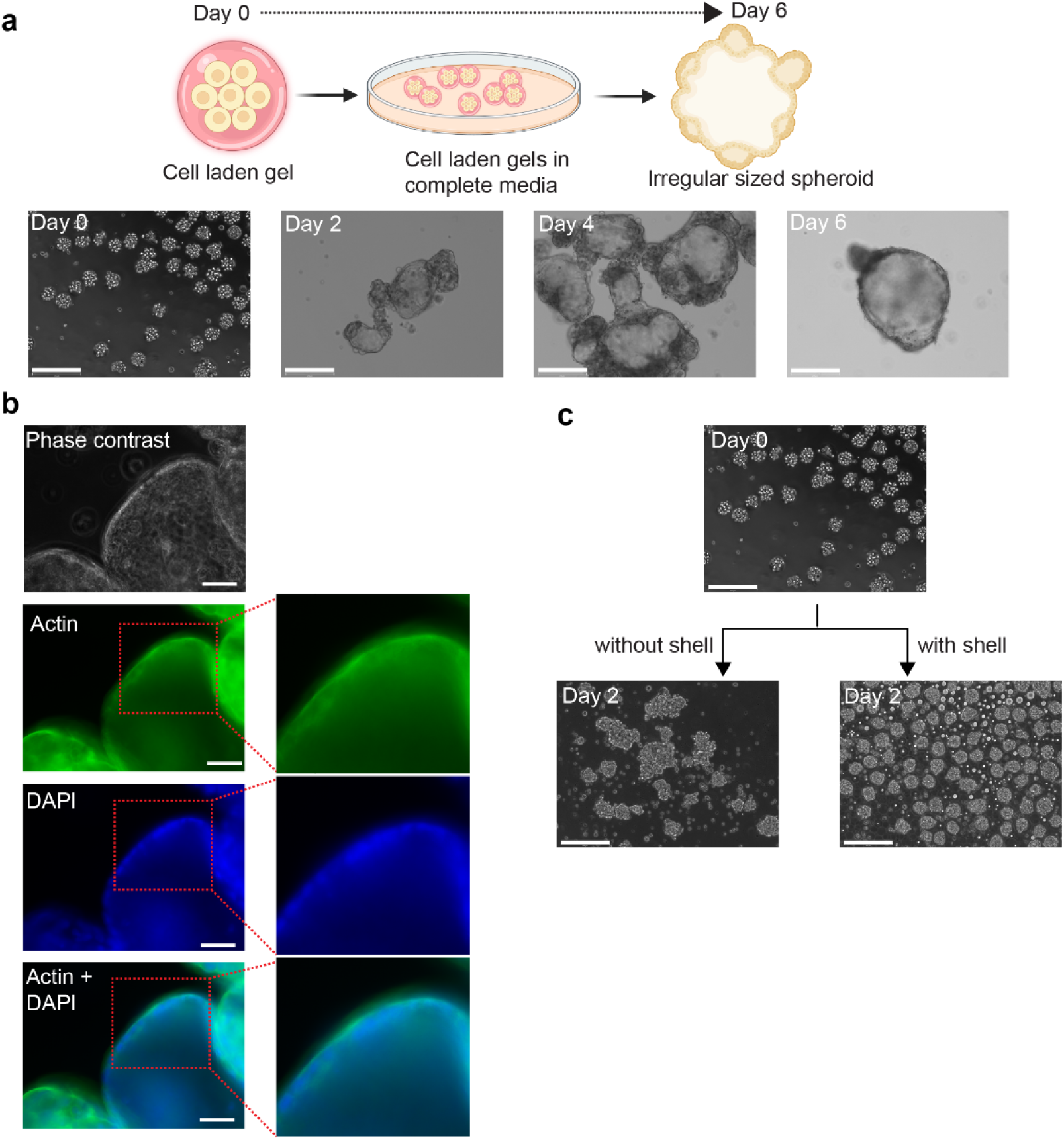
Morphological development and encapsulation effects on spheroid formation. **a,** Schematic illustration of spheroid formation in cell-laden Matrigel beads cultured over 6 days. Phase-contrast images show progressive aggregation and compaction from day 0 to day 6. Scale bars, 275 µm. **b,** Phase contrast and fluorescent images of spheroids formed by cells cultured on Matrigel beads, stained for actin (green) and nuclei (DAPI, blue), highlighting epithelial organisation in an apical-out orientation. Insets show magnified regions of apical actin enrichment. Merged images confirm cortical actin localisation surrounding nuclei. Scale bars, 150 µm. **c,** Phase-contrast images comparing spheroid culture without shell (left) and with agarose shell encapsulation (right) at day 2. Encapsulation promotes more uniform spheroid formation and prevents aggregation between spheroids. Scale bars, 275 µm.

**Supplementary Fig. 2.**
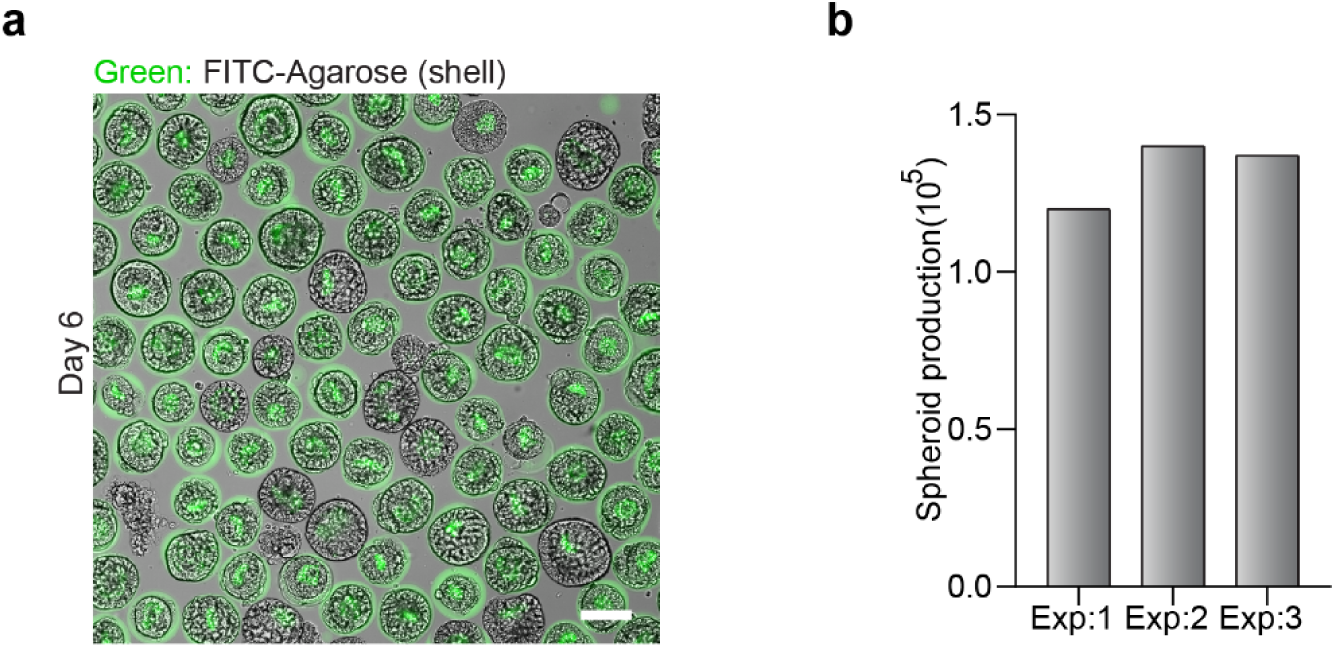
Large-scale spheroid production per experiment. a, Representative merged fluorescence and phase-contrast image of spheroids at day 6, illustrating large-scale production within a single experiment. Agarose shells are labelled with FITC (green). Scale bar, 50 µm. b, Total spheroid yield per independent experiment (Exp. 1–3). Bars represent the total number of spheroids generated per experiment, demonstrating consistent production exceeding 1 × 10⁵ spheroids per run.

**Supplementary Figure 3.**
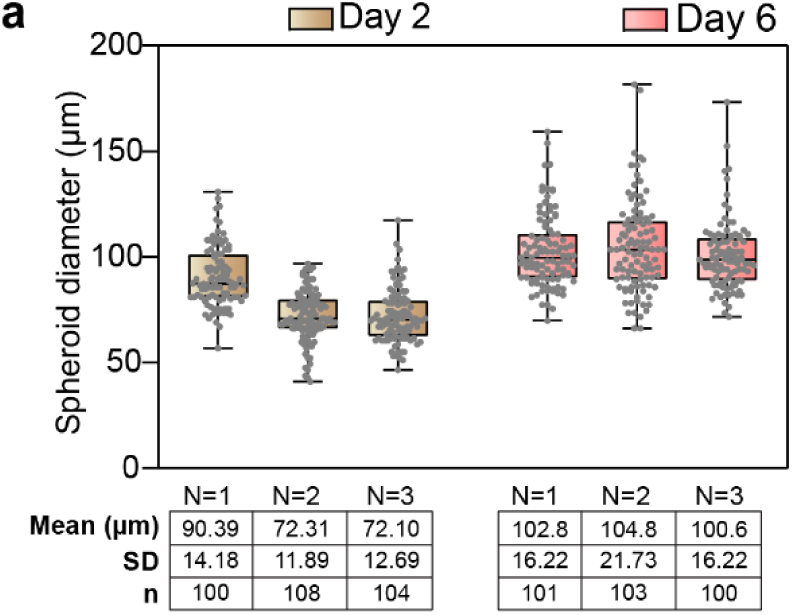
Evaluation of spheroid formation, growth, and viability in agarose microcapsules (related to Fig. 2d). a, Box plots showing spheroid diameter distributions at day 2 and day 6 across three independent experiments (N = 1–3). The table below summarises mean spheroid diameters (µm), standard deviations (s.d.), and the number of spheroids analysed (n).

**Supplementary Fig. 4.**
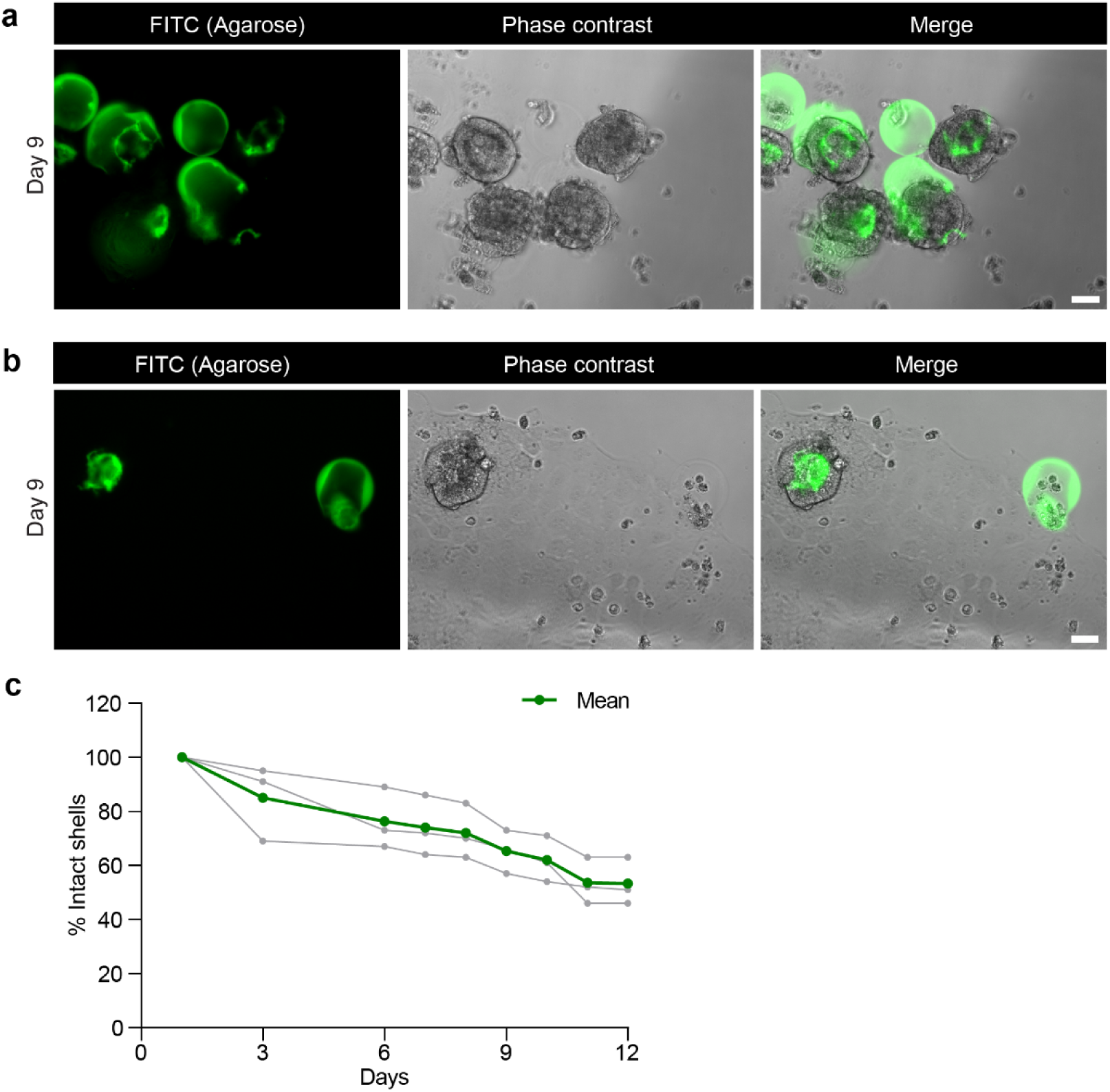
Longitudinal assessment of agarose shell integrity during extended culture. **a,** Representative fluorescent and phase-contrast images of microcapsules at day 9 showing examples of shell rupture (FITC-labelled) with spheroids breaching the capsule boundary. Scale bar, 50 µm. **b,** Representative fluorescent and phase-contrast images of spheroids at day 9 released from ruptured shells and attaching to the culture surface, resulting in 2D outgrowth. Scale bar, 50 µm. **c,** Quantification of the percentage of intact agarose shells over 12 days of culture. Grey lines indicate individual biological replicates (N=3; 100 shells each), and the green line denotes the mean, demonstrating a progressive decline in shell integrity over time.

**Supplementary Figure 5.**
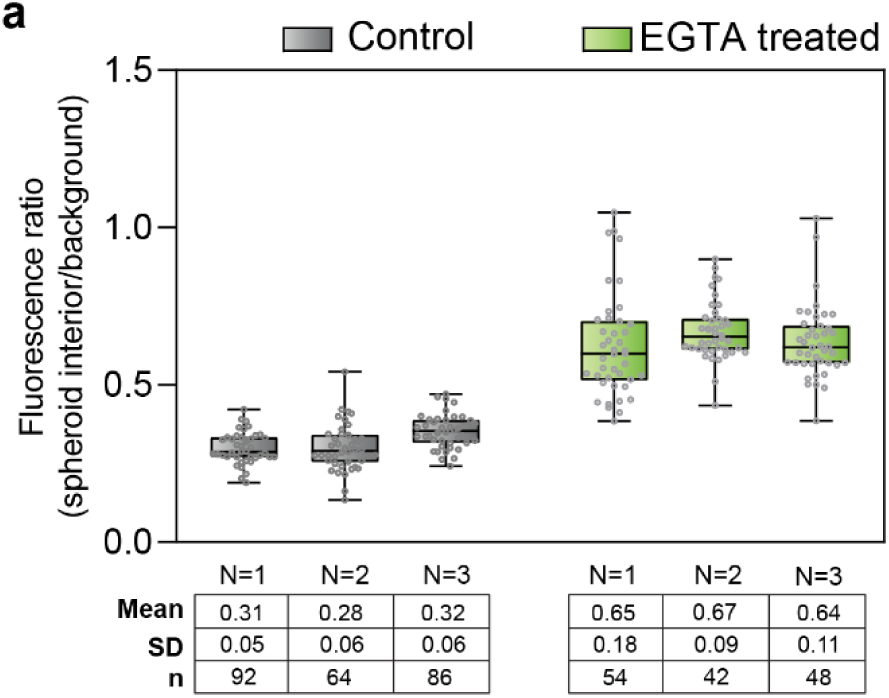
Evaluation of barrier integrity in microcapsules using FITC-dextran assay (related to Fig. 4b). a, Box plots showing fluorescence ratio of control and EGTA-treated spheroids across three independent experiments. The table below summarises mean spheroid ratio, standard deviations (s.d.), and the number of spheroids analysed (n).

**Supplementary Figure 6.**
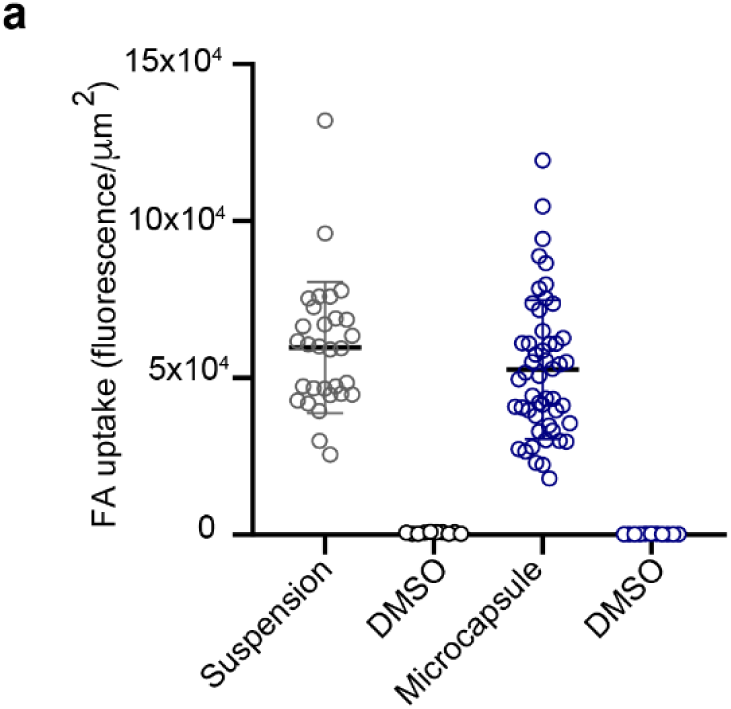
Quantification of free fatty acid (FFA) uptake in spheroids generated by suspension culture versus microcapsules (related to Fig. 5b). a, Scatter plots show C1-BODIPY-C12 fluorescence intensity normalised to spheroid area (fluorescence/µm²). Spheroids formed in microcapsules exhibited comparable fatty acid uptake to those generated by suspension culture, whereas DMSO controls showed negligible signal. Data points represent individual spheroids (microcapsules, n = 50; suspension culture, n = 10), collected across three independent biological replicates. Horizontal bars indicate mean ± s.d.

